# Microglia orchestrate neuronal activity in brain organoids

**DOI:** 10.1101/2020.12.08.416388

**Authors:** Ilkka Fagerlund, Antonios Dougalis, Anastasia Shakirzyanova, Mireia Gómez-Budia, Henna Konttinen, Sohvi Ohtonen, Fazaludeen Feroze, Marja Koskuvi, Johanna Kuusisto, Damián Hernández, Alice Pebay, Jari Koistinaho, Sarka Lehtonen, Paula Korhonen, Tarja Malm

## Abstract

Human stem cell-derived brain organoids provide a physiologically relevant *in vitro* 3D brain model for studies of neurological development that are unique to the human nervous system. Prior studies have reported protocols that support the maturation of microglia from mesodermal progenitors leading to innately developing microglia within the organoids. However, although microglia are known to support neuronal development in rodents, none of the previous studies have reported what is the impact of microglia on neuronal growth and maturation in human brain organoids. Here we show that incorporating microglial progenitors into the developing organoid supports neuronal maturation, the emergence of neurons capable of firing repetitive action potentials and the appearance of synaptic and neuronal bursting activity. Immunocompetent organoids enable experimental strategies for interrogating fundamental questions on microglial and neuronal diversity and function during human brain development.

## Introduction

Human brain organoids, generated from either embryonic or induced pluripotent stem cells, recapitulate the tissue architecture of the developing cortex to a remarkable degree of fidelity, even discriminating species differences in the development dynamics of neurons between humans and gorilla^1^(BioRxiv). Since the initial nearly parallel work by three research groups^2–4^, brain organoids have served in modelling various brain diseases^5,6^, including microcephaly^2^, glioblastoma^7,8^, Zika-virus infection^9^ and Alzheimer’s disease^10–12^.

The accumulating body of scientific reports portray brain organoids as a human 3D in vitro platform that achieves remarkable complexity, even resembling its *in-vivo* counterpart, the embryonic human brain^13–17^. The main cell types of the brain, astrocytes and neurons, spontaneously develop through ectodermal lineage in brain organoids and self-assemble to form cortical layers resembling human brain organization^18^. However, microglia, the brain’s immune cells, do not innately populate brain organoids, since they originate from mesodermal lineage, unlike other brain cells^19–21^. In addition to their roles in immunity, microglia participate in forming and molding the CNS on multiple stages of development from neurogenesis to maturation of synapses and neural networks^22–25^. During early development, microglia restrict the number of neuron progenitors arising at the subventricular zone (SVZ)^22^ and support the differentiation of neurons, astrocytes and oligodendrocytes^26,27^. The dual role of microglia is evident also later in development: Microglia support the formation of synapses through direct contact^24^ and indirectly through secretion of cytokines^28,29^. Yet, they restrict the number of neuronal connections by phagocytosing inactive synapses through the complement cascade^23,30^. Microglia also modulate the formation of neural circuits^25,31^. In addition to neurodevelopment, microglia have a distinguished role in the progression of neurodegenerative diseases^32^. Thus, the past years have seen multiple approaches to incorporate microglia into brain organoids.

The first such report showed microglia derived from induced pluripotent stem cells to migrate spontaneously from culture medium to the brain organoid. Therein, they assume their native functionality by migrating to the site of tissue injury, induced by a needle prick^33^. In an alternative approach, subtle adjustment to one of the original organoid protocols^2^, gave rise to a native population of microglia that were capable of immune response and phagocytosis of synaptic material, as shown by internalisation of post-synaptic density protein 95 (PSD95)^34^. In another report, the incorporation of mesodermal progenitors introduced vasculature into the organoid, and remarkably, also microglia^35^. Finally, a very recent report showed that transplantation of human primary prenatal microglia to brain organoids gave rise to synchronous burst activity^36^ (BioRxiv). Synchronous burst has also been reported in the absence of microglia, though in much older organoids, and in conjunction with emerging inhibitory interneurons^16^.

Here we carry out detailed single-neuron interrogation of electrophysiological functions that arise in brain organoids in response to incorporation of microglia. We show for the first time that in organoids with microglia, neurons respond to a depolarising stimulus with sustained repetitive firing, while neurons without microglia are more prone to adaptation. Remarkably, neurons co-maturing with microglia showed a greater diversity in the expression of distinct neuronal currents, including a prominent post-inhibitory low-threshold current leading to post-inhibitory spiking*, an action* attributed to T-type calcium channels, which have been implicated in a variety of functions, including synaptic plasticity and neuronal differentiation^37–39^. We also found that excitatory post-synaptic currents, evidence of synaptically connected neurons, were present only in neurons from organoids with microglia. Finally, targeted single-cell and multielectrode array recordings revealed regular spontaneous neuronal bursting activity only in organoids with microglia. Taken together these data suggest that multiple aspects of single-neuron functional characteristics are strongly impacted by microglia, indicating the emergence of microglia-dependant sub-population of neurons in cerebral organoids. Our work herein consolidates brain organoids as a tool to study the roles of microglia in brain development, in ways not accessible *in-vivo.*

## Results

### 1. Erythromyeloid progenitors migrate into brain organoids and mature into microglia-like cells

Given that microglia migrate into brain early during embryogenesis, we hypothesized that erythromyeloid progenitors (EMPs) would migrate spontaneously to brain organoids at an early state, as had earlier been shown for more mature induced pluripotent stem cell (iPSC)-derived microglia^33^. To test this idea, we differentiated iPSC-derived erythromyeloid progenitors (EMPs) according to previously published microglial differentiation protocol^40^. We incorporated them in day 30 cerebral brain organoids which we generated from the same iPSCs as previously published^2^. A Schematic illustration of merging these protocols is described in Fig. 1a. Remarkably, the incorporated EMPs establish a microglia-like population and colonise the organoid, as presented in Fig. 1b. The robustness of differentiation was confirmed by performing differentiation with and healthy iPSC lines (**Table**). Immunohistochemical stainings against the EMP marker CD41 revealed the presence of the EMP population suspended in matrigel adjacent to the organoid tissue still five days after the incorporation, at day 35 (Fig. 1c). Cells positive for the microglia marker Iba1 and negative for CD41 had begun to infiltrate the nascent cortical loops, as identified by their tissue architecture shown in immunohistochemistry (IHC) staining against TBR2 positive intermediate progenitors and DCX positive young neurons (Fig. 1d). Moreover, Iba1+ cells attained less spherical morphology upon arriving at the organoid tissue, and as the organoids matured, the Iba1+ cells portrayed a spectrum of morphologies (Fig. 1e, day 120). The organoids showed radial maturation of the cortical plate (Fig. 1f, day 120), as shown previously^2,18^ and the Iba1+ cells exhibited tangling with neuronal processes (Fig. 1g, day 66). Also, in accordance with earlier work, we observed minor native populations of microglia in organoids without incorporated microglia progenitors^34^. However, counting the Iba1+ cells in day 120 organoids with and without incorporated microglia, abbreviated here (+)ORG and (-)ORG, respectively, showed significantly more microglia in the (+)ORG group (S.Fig. 1a)

**Figure 1.**
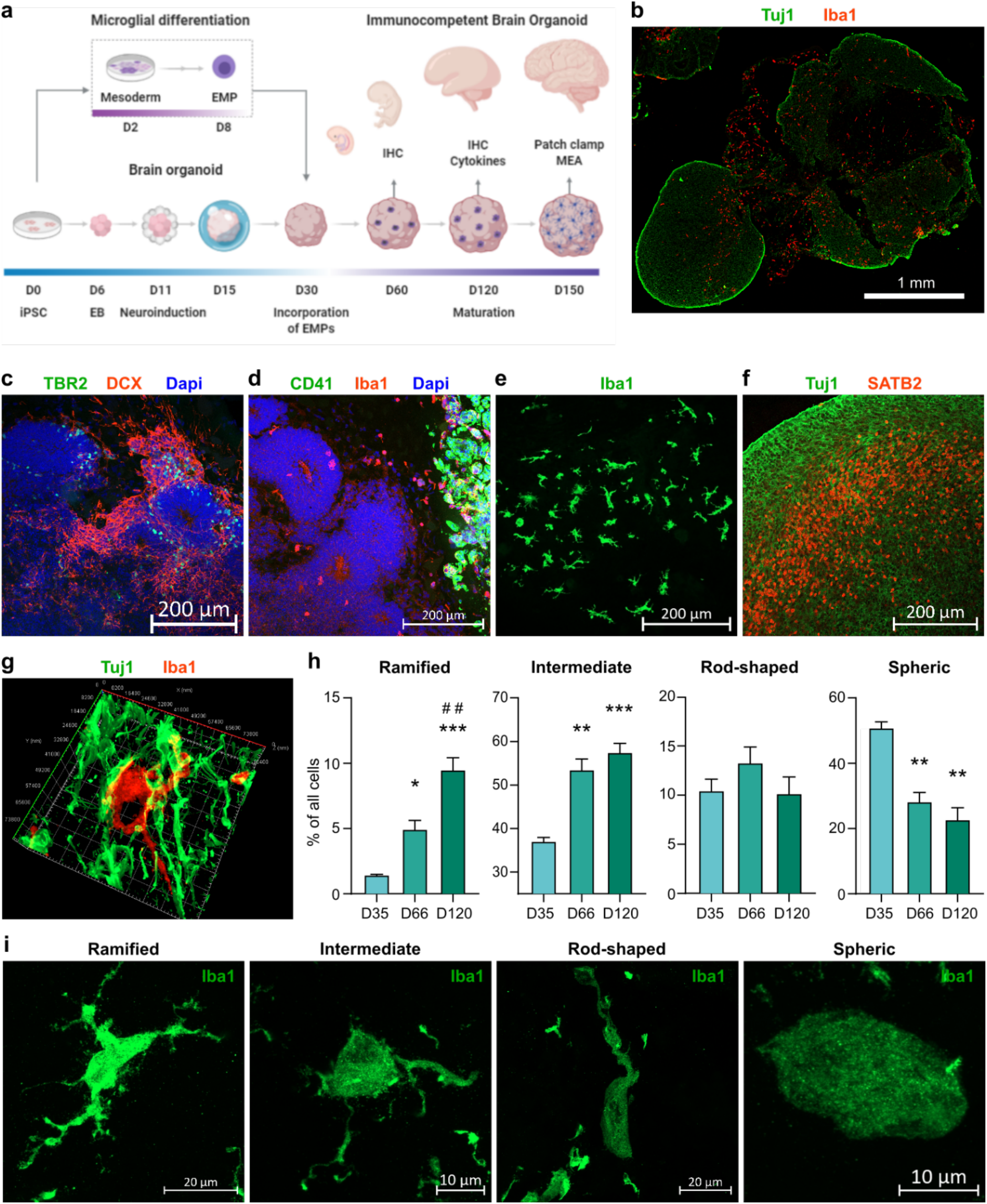
Erythromyeloid progenitors migrate to establish to a microglia-like population in cerebral organoids. **a** Schematic drawing depicting the introduction of EMPs to ORGs and subsequent experiments. **b** Immunohistological staining of Iba1+ cells colonizing a Tuj1 stained cORG at day 120. **c** and **d**, immunohistological stainings of adjacent sections of an ORG at day 35, one day after incorporation of EMPs. TBR2+ and DCX+ cells show the formation of cortical loops, while CD41 shows the locus of incorporation and Iba1, the cells migrating from the locus to the vicinity of the nascent cortical loops. **e** A population of Iba1+ cells in taking different morphologies. **f** Maturation of a cortical loop, as shown by immunostaining of SATB2 in a day 120 organoid. **g** A 3D reconstruction of an Iba1+ cell and its processes closely intertwined with Tuj1+ neurons in a day 66 organoid. **h** Detection and quantification of different morphology types of Iba1+ cells in organoids on days 35, 66 and 120, using the artificial intelligence platform by Aiforia. The Y-axis presents the portion of the respective morphology type in all the detected Iba1+ cells. The bars represent the mean (+/- SEM) percentage of three organoids for the day 35 group (one batch), six for day 66 (one batch) and nine for day 120 (two batches), Tukey test, significance ***p < 0.001, **p < 0.01, *p < 0.05 compared to D35, # indicate similarly significance compared to D66; **i** Representative orthogonal projection images of all the quantified morphology types, imaged from a single section of a day 120 organoid. Note the higher Iba1 intensity in the ramified cell, as well in the process residues, neighbouring the rod-shaped cell, orphaned from their respective cell somas during cryosectioning.

As the microglia take increasingly complex morphologies during brain development *in-vivo^41^,* and given the breadth of variation in the morphologies of Iba1+ cells even within a single organoid, we sought to answer if the relative portions of certain particular morphologies within the Iba1+ cell population changes as the organoid matures. For this, we employed an artificial intelligence (AI) platform offered by Aiforia and trained the convolutional neural network-based algorithm to annotate four different morphology types (Fig. 1i) from images of immunohistological stainings against Iba1 in organoid cryosections. We quantified the numbers of ramified, intermediate, rod-shaped and spheric cells in organoid sections at days 35, 66 and 120 into the organoid culture protocol (Fig. 1h). On day 35, the Iba1+ populations consisted mainly of cells detected as spheric and intermediate. The major changes on day 66 shifted to more ramified and intermediate morphologies and decreased spheric cells. On day 120, the ramified cells were further enriched within the Iba1+ population, while the portion of other types did not significantly change from day 66. In parallel to the AI-based quantification, we performed skeletal analysis using ImageJ^42,43^, entailing manual annotation of the cells and yielding per-organoid average counts of branches, junctions, endpoints and triple points (S.Fig. 1b). In agreement with the AI-based detection results, the average complexity of cell morphology increased from day 35 to 66. However, the complexity decreased from day 66 to day 120. Combining the results from the AI-based classification and skeletal analysis, suggests that indeed, microglia-like cells attain more complex morphologies as they mature within the organoid environment. However, while the average morphology becomes less ramified from day 66 to day 120, as shown by skeletal analysis, the portion of the ramified subset increases, as demonstrated by AI-based detection.

### 2. Microglia interact with synapses but do not make the organoids immunologically responsive

IHC staining against Iba1 and B-tubulin showed the microglia to be intensively intertwined within neurons (Fig. 1g), prompting us to further investigate physical interaction between the microglia and synaptic material. Indeed, double staining for Iba1 and post-synaptic density protein95 (PSD95), and Iba1 and the presynaptic synaptophysin 1 (Syn1) revealed instances of synaptic material being inserted in an open pocket on the surface of the microglia (Fig. 3a-b). Orthogonal dissection images revealed Syn1-stained material to fill the entire pocket, while PSD95 filled only part of the pocket, suggesting a greater part of the post-synapse to be seated in than what was revealed by the PSD95 staining. Also, for both Syn1 and PSD95, we observed partial co-localisation with Iba1 within the pocket. For Syn1, we also observed instances of non-pocketed co-localisation with a microglial process. Taken together, the images compose immunohistology evidence of the incorporated microglia physically interacting with both pre- and post-synapses.

Next, to analyse if the interaction between microglia and synapses was reflected in the numbers of PSD95 puncta, we quantified the number of PSD95 puncta in the (+)/(-) ORGs (S.Fig. 1c). Interestingly, in each prominence threshold, the average PSD95 puncta counts were slightly increased in the (-)ORG group, yet the difference failed to reach statistical significance. Also, we observed two distinct spatial patterns of PSD95 expression (S.Fig. 1d); one, where the puncta are scattered across a region in the organoid section, and the other, where the puncta are not scattered but instead localised in cell soma. Most organoids presented areas with both spatial distribution patterns of PSD95, and we detected no apparent differences in the prevalence of cells with soma localised PSD95 between the (+)ORG and (-)ORG groups.

Since microglia are the immune cells of the brain, we evaluated whether incorporation of microglia would evoke functional pro-inflammatory responses lipopolysaccharide (LPS). We measured the concentrations of the pro-inflammatory cytokines TNF, RANTES, IL10, IL6, IL1a, IL1b, IL8 and MCP1 from the culture medium after 24 hours of exposure to 20ng/ml LPS. Only IL8 and MCP1 were secreted in detectable concentrations in the culture medium, with no significant difference between LPS and vehicle exposure between the (-)ORG and (+)ORG groups (Fig. 2d). However, the basal secretion of IL8 and MCP1 in vehicle-treated organoids showed that the (+)ORG medium had significantly more MCP1 compared to the (-)ORG medium (Fig. 2c). LPS exposure did not cause evident changes in Iba1+ cellular morphologies as analysed by the AI-based detection (Fig. 2e) or in the average complexity of the cells, as evaluated by skeletal analysis (S.Fig. 1e). Taken together, the results indicate that while the incorporated microglia did increase basal secretion of the pro-inflammatory MCP1, our LPS exposure paradigm failed to induce significant cytokine secretion or microglial morphological alterations in the organoids.

**Figure 2.**
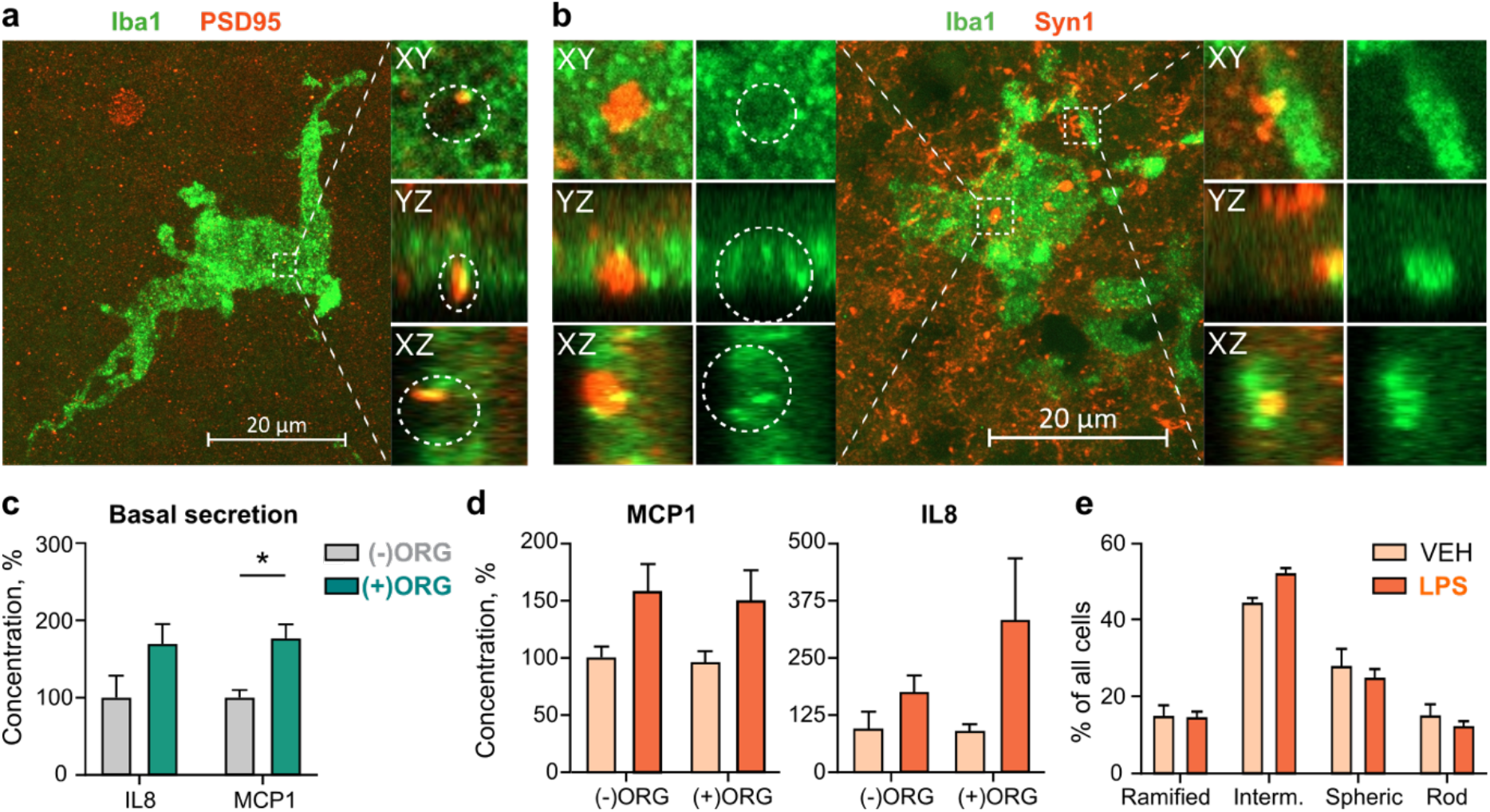
Incorporated microglia interact with synapses but do not show immune response. **a** An orthogonal projection and orthogonal dissections of PSD95+ post-synaptic material embedded within a pocket on the surface of an Iba1+ cell. **b** Instances of Syn1+ presynaptic material embedded within a pocket on the soma of Iba1+ cell and partially internalised by process of the cell. **c** Basal cytokine secretion from the vehicle treated organoids show a significant increase in the (+)ORG group for MCP1 but not for IL8. Data represent the normalised values from 8-10 organoids. The measured cytokine concentrations were normalised to the mean of the (-)ORG vehicle group for for each cell line, one batch per cell line. The normalised values of each cell line were pooled together to test for group differences. **d** Normalized (as in **c**) concentrations of the proinflammatory cytokines MCP1 and IL8 measured from the culture medium of organoids with and without incorporated microglia after 24-hour exposure to LPS or vehicle. The bars represent normalised means (+/- SEM) from 9-15 organoids at day 77 and 120 and three cells lines (Mad6, Bioni-010-C2 and Bioni-010-C4) per group, one batch per cell line. **e** Detection and quantification of portions of different morphology types of Iba1+ cells after 24 hours of LPS or vehicle exposure. Bars represent means (+/- SEM) of nine organoids for vehicle and ten for LPS, MBE2968, two batches. Two-way-ANOVA, significance *p < 0.05.

### 3. Microglia expedite neuronal maturation in cerebral organoids

To assess whether enrichment of organoids with microglia has the capacity to alter single neuronal properties in the developing organoids, we performed whole-cell recordings in acute organoid slices at days 107-120 and days 150-165. The recordings were carried out under voltage-clamp conditions to measure the maximal sodium & potassium (holding voltage – 70 mV, test voltage −20 mV and +10 mV, respectively) and passive leak current density (holding voltage – 50 mV, test voltage −120 mV) as markers of neuronal maturation and stage of development. Currents recorded were normalised to cell size and expressed as pA current per pF of cell capacitance. We detected a statistically larger potassium current density in (+)ORG at the early time point and a significantly larger mean sodium current density in neurons in (+)ORG at both recorded time points (Fig. 3a, b). No changes were seen in the background mean maximal leak current density between the two groups (Fig. 3a, b) or basal electrophysiological parameters of the neurons (input resistance, resting membrane potential or membrane time constant, data not shown). We found that the increased sodium current density translated functionally to a larger prevalence of repetitive action potential (AP) firing neurons in (+)ORG (Fig. 3e) and a larger prevalence of neurons firing non-adaptively during the current steps (Fig. 3f). Also, neurons from (+)ORG were more likely to express active ionic currents and, in particular, a presumed low-threshold calcium current often leading to a low threshold potential-spike (LTS), following release from a prolonged hyperpolarisation (Fig. 3g, h). Furthermore, we only detected spontaneous excitatory post-synaptic currents (sEPSCs) under voltage-clamp in neurons in (+)ORG but not in (-)ORG (Fig. 3i, k).

**Figure 3.**
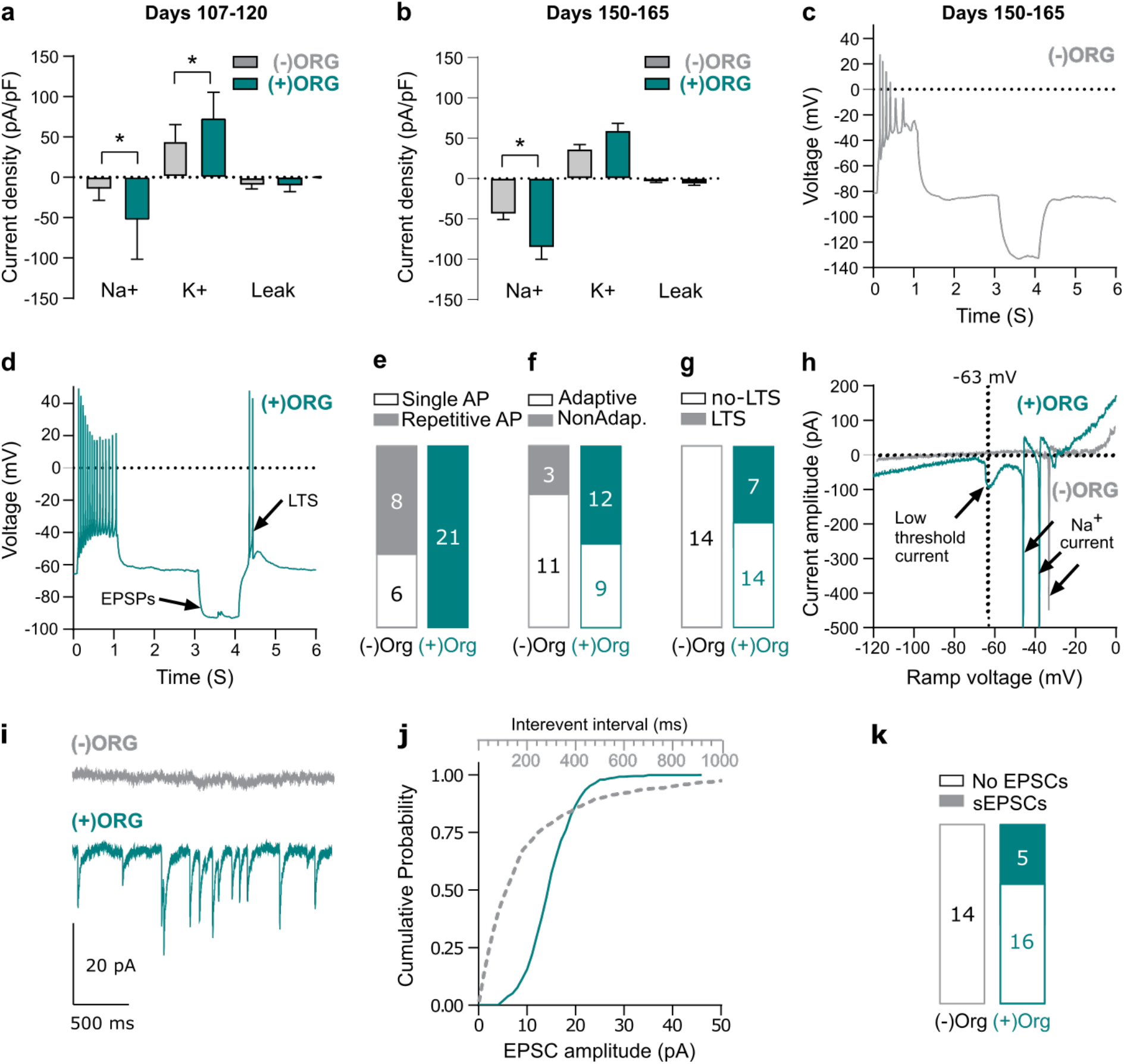
Microglia empower neuronal maturation and synaptic signalling in (+)ORGs. **a** Maximal sodium, potassium, and leak, current density in 107-120 day old organoids. N= 8 vs 9 neurons (mean ± SEM; Mann-Whitney unpaired test, *p<0.05, 4 to 5 organoids per group from two independent batches). **b** Maximal sodium, potassium and leak current density in 150-165 day old organoid. N=14 vs 21 neurons (mean ± SEM; Mann-Whitney unpaired test, *p<0.05, 6 to 9 organoids per group, from three independent batches. **c** Electrophysiological traces recorded in current-clamp mode showing the responses of an (-)ORG neuron to positive and negative current injections (± 12.5 pA, 1s). Note the AP amplitude, complete cessation of firing at the end of the depolarising step and the pure passive responses during and after the hyperpolarising step. **d** Electrophysiological traces recorded in currentclamp mode showing the responses of an (+)ORG neuron to positive and negative current injections (±20 pA, 1s). Note the large AP amplitude, the sustained firing at the end of the depolarising step and the low threshold spike after the end of the hyperpolarising step. **e** Summary results for single AP and repetitive AP firing neurons recorded in (-)ORG and (+)ORG slices. **f** Summary results for spike frequency adaptation during the 1 s depolarizing step recorded in (-)ORG and (+)ORG neurons. **g** Summary results for the presence of post-hyperpolarisation low threshold spike-potential (LTS/LTP). **h** Electrophysiological traces recorded with a voltage ramp (speed 0.14 V/s) taken from neurons shown in **c** & **d**, showing an inward current developing in the subthreshold range, which is responsible for the LTS responses recorded in the (+)ORG neuron. Note the complete absence of current nonlinearities in the (-)ORG neuron in subthreshold voltage potentials. **i** Electrophysiological traces of spontaneous excitatory postsynaptic currents (sEPSCs) recorded under voltage clamp (holding potential of −70 mV) from an (-)ORG and (+)ORG neuron. Note the complete absence of sEPSCs in the (-)ORG neuron. **j** Cumulative distribution curves for inter-event interval and amplitude of sEPSCs detected on the (+)ORG neuron presented in **h**. **k** Summary results for the presence of sEPSCs in (-)ORG and (+)ORG neurons. Non-parametric Mann-Whitney unpaired test, significance *p < 0.05.

### 4. Microglia mediate formation of spontaneous bursting activity in cerebral organoids

Next, we looked if microglia would boost the emergence of network activity, previously reported in organoids^13^. 3D-MEA electrophysiological recordings were undertaken from acute slices of day 150–165 organoids to examine the developing characteristics of the neuronal network properties. We only detected a single active electrode track with spontaneously occurring multiunit activity from (-)ORG slices, which was in striking contrast to the (+)ORG slices where we detected 24 spontaneously active electrode tracks under basal conditions (Fig. 4a, b). (+)ORG slices exhibited activity patterns in active electrode tracks ranging from irregular firing to regular bursting (Fig. 4b-d). Such bursting activity could be often discriminated successfully, through principal component analysis, into single unit activity (SUA; Fig. 4c) from presumed single neuron firing, but each unit appeared relatively asynchronous in bursting phase to others and thus population spiking plots remained flat (Fig. 4d). and did not follow any oscillatory activity with typical high amplitude peak spiking-low amplitude silent pause features expected from synchronously bursting networks as seen before by others^13^. The differences in the prevalence of activity and functional characteristics were not secondary to slice-MEA placement or viability of the neurons as we could reverse the silent electrode tracks of the (-)ORG slices to active by exciting the neurons with the application of NMDA, resulting in the emergence of brief bursting activity (Fig. 4h, a). Whole-cell recordings from organoids that have undergone MEA recordings confirmed the presence of bursting neurons from resting (0 pA) or near resting membrane potential (from +2 to +10 pA current injection, Fig. 4g) in slices from (+)ORG but rarely from slices of (-)ORG (Fig. 4f).

**Figure 4.**
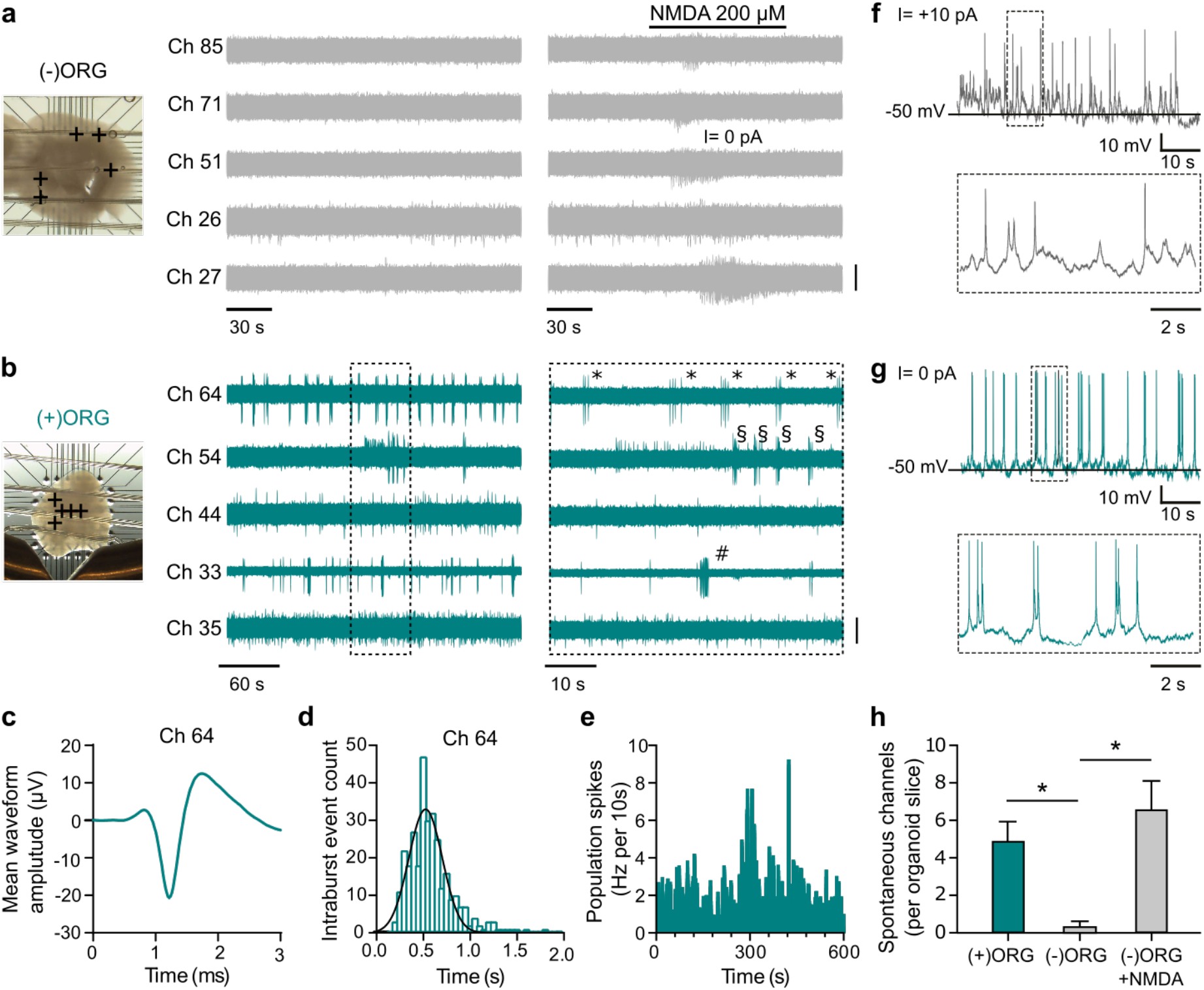
Microglia drive neuronal bursting and network activity in (+)ORGs. **a** Image of an (-)ORG slice recorded with a 3D 8×8 MEA (left) and raw voltage recordings of spiking activity from five representative electrodes (depicted as + symbols on image) before and after NMDA perfusion (middle and right panels). Only channel 26 exhibited activity in baseline conditions (one active electrode detected from 4 organoid slices), while NMDA perfusion effectively and quickly elicited a brief burst of activity on all five channels (bar, 20 μV). **b** As described above in **a** but for a (+)ORG slice. The regular bursting activity of 3-7 spikes was detected in three electrodes (33, 54, 64) (twenty-four active electrodes detected from 5 -organoid slices). Symbols mark individual bursts in each different channel (bar, 20 μV, except for Ch 33, 40 μV). **c** Mean spike waveform after spike sorting (255 spikes) using principal component analysis from raw signal depicted in **b** from channel 64. A single tight cluster with a well-defined AP waveform was detected consistent with the recording being from a single neuron (single unit activity, SUA). **d** Intra-burst spike interval histogram (50 ms bins) for SUA from channel 64 presented in **b, c**. Ten of total 24 active electrode tracks detected from 5 (+)ORG slice-organoids were bursting. The single Gaussian function fitted to the data is centered at 524 ± 178 ms (mean ± SEM) while the mean inter-burst interval for this neuron was 10.7 ± 4.2 s (mean ± SEM). **e** Population spike activity from all the active electrodes the (+)ORG slice presented in **b** (in 100 ms bins). Although individual electrodes exhibited bursting activity, this activity was largely asynchronous between electrodes, as seen by the typical flattened population spiking response over the time course of 10 minutes. **f** Electrophysiological traces from whole-cell recording from a (-)ORG neuron exhibiting a current-induced (+10 pA) low amplitude, short term bursting behavior. **g** Electrophysiological traces from whole-cell recording from a (+)ORG neuron exhibiting high amplitude, robust, regular bursting behavior from its resting membrane potential. **h** Bar chart comparison of active electrode tracks recorded per organoid slice. (+)ORG slices exhibited a statistically significant larger number of detected active electrode tracks compared to (-)ORG slices (. This lack of spontaneous activity could be reversed by perfusion of NMDA that caused the emergence of transient activity on previously silent electrodes recorded from (-)ORG slices (26 active electrodes from 4 organoid slices, n=4 (-)ORGs and n=5 (+)ORGs from two independent batches).

These data suggest that neurons in (-)ORG are spontaneously inactive in extracellular recordings and do not have the capacity to fire coherent bursting patterns. In contrast, neurons in (+)ORG exhibit clear signs of regular and robust single-cell bursting characteristics without having yet developed concrete network features of synchronised population activity. The larger sodium-potassium current density, the more consistent firing & bursting characteristics and the presence of greater ion channel diversity and of synaptic activity in single neurons of microglia enriched organoids examined here suggest that microglial enrichment has affected the development and the fundamental aspects of maturation of single-cell and neuronal network properties in organoids.

## Discussion

Here we show that the presence of microglia drives neuronal maturation in cerebral organoids. To our knowledge, this is the first study providing evidence that the presence of microglia dictates the appearance of single neurons with more mature functional properties as compared to neurons that mature without direct contact to microglia.

Previous landmark papers have reported successful introduction of microglia to brain organoids, each with very different approaches. Abud et al. (2017)^44^ showed incorporation of mature iPCS derived microglia to mature organoids and included MCSF1, IL34 and TGFB in the culture medium to feed the microglia their most crucial growth factors. Soon followed, Ormel et al. (2018)^34^ adjusted the original forebrain organoid protocol^2^ to allow native EMPs to survive and differentiate to microglia. Our approach fits between the two ends of the spectrum. We show that exogenous EMPs migrate into young brain organoids, shed away the EMP marker CD41 and mature to microglia-like cells, while completely relying on the growth factors and cues offered by the organoid tissue environment. Analogically to work by Ormel et al (2018)^34^ and with previous results from *in-vivo*^41^, we showed the incorporated microglia to attain more ramified morphology, as the organoid matured, while displaying considerable morphological differences within a single organoid, in agreement with their *in-vivo* counterparts in the developing brain^45,46^.

At the time of incorporation, the microglia progenitors expressed CD41, a marker of primitive hematopoietic precursors, as is likewise expressed by microglia progenitors when they migrate into the developing brain^47,48^. At the time of incorporation, the cells are still able to proliferate^40^ and thus, the yielded number of microglia in the organoids may vary. As our incorporated microglia rely completely on the organoid environment for maturation and survival, it may be that some organoids have a more supportive environment than others, giving rise to the differences in microglia counts we observed. Analogically, some areas within the organoid may have chemotactic cues that lure microglia^49^. And further complicating the scenario, these cues may originate from developing neurons or the emerging necrotic areas, as both phenomena occur in parallel during maturation of the organoids^2^.

However, surprising and exciting changes in microglia population dynamics occurred when moving from day 66 to day 120. While the average cell morphology reverted to a less complex shape, the population of ramified cells increased from 5% of all microglia on day 66 to 10% on day 120. Thus, as the portion of microglia with more mature morphology increased, the great majority of cells appeared less mature or more activated. Indeed, we speculate that the observed morphological degeneration at day 120 arose due to the formation of the necrotic core in the older organoids, with the ramified population existing in niches less exposed to cues from the necrotic core. Microglia are known to respond to necrosis *in-vivo^50^.* Thus, while aiming to measure morphology change due to maturation, we may have, in fact, detected the morphology change related to the activation of microglia^50^. If so, it may have offered an unintended but welcome indication of functional response and portraying the incorporated microglia as cells that are actively surveying and responding to environmental cues, akin to their *in-vivo* counterparts^49^.

The incorporated microglia induced minor but significant basal secretion of the pro-inflammatory MCP1, suggesting the incorporated microglia hold some immunological functionality. However, we could not evoke further significant proinflammatory response or changes in microglial morphology in organoids exposed to LPS. This may be due to the age of the organoids. If, indeed, the necrotic core was already activating the microglia in the day 120 organoids, then any further activation by LPS may have been too incremental to be caught in our measurements. Further experiments with younger organoids may be justified, as previous work has shown LPS evoked secretion of pro-inflammatory cytokines in far younger organoids, at around 7 weeks^34,51^.

Physical interaction between microglia and synapses in neurons has been convincingly shown *in-vivo*^24^. Ormel et al (2018)^34^ reported the first such proof in brain organoids, showing co-localisation of post-synaptic elements and microglia. Likewise, our study shows co-localisation, which may indicate synaptic pruning, an essential developmental step to eliminate excessive and little used synapses^23^. In addition, we revealed fascinating instances of synaptic pocketing, where microglia isolate a synaptic element in an open cavity, which could be a step prior to phagocytosis. However, the more mature phenotype of neurons in the (+)ORGs, as caught in our electrophysiological recordings, encourages us to speculate that, rather than pruning, the direct intercellular contact may mediate maturation of synapses, as has been described *in-vivo*^24^. Importantly, our data also show that the total number of synapses, as reflected by the PSD95 puncta counts, did not drastically differ between the (-) and (+)ORGs, indicating that even with the possible inflammatory priming by the necrotic core, the incorporated microglia did not extensively phagocytose synapses, as has been shown in neurodegenerative diseases, where microglia shift to inflammatory phenotype^52^.

Inspired by the close interaction with microglia and synapses, we demonstrated mature and diverse neuronal phenotypes to emerge in organoids with microglia. We recorded spontaneous neural activity using multi-electrode arrays in neuroimmune and control organoids at day 150 and found increased prevalence of single neurons with oscillatory bursts in immunocompetent organoids compared to control consistent with the presumed role of microglia in network formation^29,53^. Our data suggest that neurons from (-)ORG are inactive in extracellular recordings and do not have the capacity to fire coherent bursting patterns. In contrast, neurons from (+)ORG exhibit clear signs of strong and regular single-cell bursting characteristics without, however, having developed yet concrete network features such synchronised population activity^16^. The larger sodium-potassium current density, the more consistent firing & bursting characteristics and the presence of greater ion channel diversity and of synaptic activity in single neurons of microglia enriched organoids examined here suggest that microglial enrichment has affected the development and the fundamental aspects of maturation of single-cell and network properties of neurons in the organoids. These data combined suggest that sodium current properties, that are taken as a sign of neuronal maturation^54^ are more developed in the neuroimmune organoid neurons, unlike the stripped-down nature of the non-neuroimmune organoids that lack specific development of these epitomes of neuronal function. Thus, microglia orchestrate brain development via modulation of maturation of spontaneous oscillatory network properties by virtue of interacting with the developing synapses.

Although our study, being the first of the kind to describe how microglia lead to single-cell neuronal maturation, our model also suffers from some limitations. The model itself, being based on the Lancaster protocol, yields quite variable organoid structures^55^. Even in microglia containing organoids, the neurons remained immature and failed to show synchronous network activity. Thus, employing protocols shown to give more mature electrophysiological functions^13,56^ yield more mature neuronal spiking properties and the ability to investigate network activity.

A plethora of knowledge on the role of microglia in neurodevelopment comes from mice, with only speculations to account for the corresponding events in humans. Addition of microglia into cORGs makes it a radically more complete platform to study human brain development in health and, especially, in disease. The immunocompetent cORG model that we have developed serves as a relevant human *in-vitro* model applicable to neurodevelopment disorders linked to maternal immune activation during pregnancy, such as autism spectrum disorders, schizophrenia and epilepsy and will help in revealing novel therapeutic targets for these diseases.

## Methods

### Human iPSC lines and ethical considerations

Four iPSC lines were used and their phenotype and origin are presented in Table 1. Three of the iPSC lines – MBE2968, Bioni010-C2 and Bioni010-C4 were previously characterized^40,57,58^. The Mad6 line was generated from fibroblasts (skin biopsy and subsequent work under Research Ethics of Northern Savo Hospital district (license no. 123/2016)) that were first expanded in Iscove’s DMEM media (Thermo Fisher Scientific) containing 20% fetal bovine serum, 1% Penicillin-Streptomycin and 1% nonessential amino acids. For reprogramming, the cells at 90% confluency were transduced using CytoTune™-iPS 2.0 Sendai Reprogramming Kit (Thermo Fisher Scientific). The media was changed 24 h after transduction and then daily. At day 6, fibroblast culture medium was replaced with Essential 6 Medium (E6, Thermo Fisher Scientific) supplemented with 100 ng/ml basic fibroblast growth factor (bFGF). The cells were re-plated on the next day to 6-well matrigel-coated plates with a density of 60 000 cells/well. The individual colonies were picked to 24-well matrigel-coated plate containing Essential 8 Medium (E8, Thermo Fisher Scientific) between days 17-28 and passaged with 0.5 mM EDTA weekly. One week later, three colonies were selected and expanded on 6-well plates with daily media changes. For RT-qPCR, (S. Fig. 1d), total RNA was extracted (RNeasy Mini kit, Qiagen), cDNA synthesized using Maxima reverse transcriptase (Thermo Fisher Scientific) and Taqman probes (Table 3). For IHC(S. Fig. 2 a), the iPCS colonies were fixed (4% paraformaldehyde, 20 minat RT, permeabilized with 0.2% Triton X-100 (in case of Nanog and Oct-4), blocked in 5% normal goat serum one hour in RT and incubated with the primary antibodies (Table 3) O/N at 4°C, followed by secondary antibody incubation one hour at RT. Images were taken at Zeiss Axio microscope, scale bar 100 μm (S.Fig 2a). For embryoid body (EB) generation, iPSCs colonies were detached by a scalpel to ultra low-adherent dishes (Corning) and cultured in DMEM media (Thermo Fisher Scientific) supplemented with 20% serum replacement and 1% Penicillin-Streptomycin. The EBs were plated onto Matrigel-coated 24- well plates and left to differentiate for two weeks. Expression of markers for all three embryonic layers was performed by flow cytometry (CytoFlexS, Beckman Coulter, Indianapolis, USA). The EBs were dissociated using accutase (4 min, at 37°C followed by trituration into single cell suspension, centrifuged 300 g 5 min, resuspended and filtered through 35 μm strainer. Cell viability was assessed by Zombie Aqua (Biolegend), cells fixed in 1% formaldehyde, permeabilised in 0.2% TritonX+ 5% FBS, and incubated with fluorophore-conjugated antibodies (Table 3) 30 min at RT.

**Table 1.** Human induced pluripotent stem cell lines and their genders, phenotype, and origin used in this study.

| Line | Sex | Health status | Age years | APO E type | Sample origin | Reprogramming method | Karyotype | Reference | Used for experiments |
| --- | --- | --- | --- | --- | --- | --- | --- | --- | --- |
| MBE2968 c1 | F | Healthy | 65 | ε3/ε3 | Skin biopsy | Episomal nucleofection | 46XX<br>Normal | 40 | Fig 1, 3 & 4, S.Fig a-c |
| Mad6 | M | Healthy | 63 |  | Skin biopsy | Sendai virus | 46XY<br>Normal |  | Fig 2, S.Fig. d & e |
| BIONI010-C-2* | M | Healthy | 15-19 | ε3/ε3 | Skin biopsy | Non-integrating episomal | 46XY<br>Normal |  | Fig 2 c-e |
| BIONI010-C-4* | M | Healthy | 15-19 | ε4/ε4 | Skin biopsy | Non-integrating episomal | 46XY<br>Normal |  | Fig 2 c-e |

\*Both lines derived from the parental BIONI010-C, only the ApoE allele differs

Karyotyoing (S. Fig. 1c) performed at Synlab oy (Finland) Line MBE2968 c1, was previously reported^40^ and the BIONi010-C-2 and BIONi010-C-4 were purchased from the European bank for Induced Pluripotent Stem cells (EBiSC) and produced and characterised by Bioneer (Copenhagen, Denmark). Mad6 line was karyotyped at the Yhtyneet Medix laboratoriot, Finland (http://www.yml.fi/).

### iPSCs culture

iPSCs were maintained in serum- and feeder-free conditions in Essential 8 Medium (E8, #A15169-01 Gibco) supplemented with 0.5% penicillin-streptomycin (P/S, #15140122 Gibco) on growth factor reduced Matrigel (Corning) at 5% CO2 and 37°C. Medium was completely changed every 24 hours. Twice a week, when cultures reached 80% confluency, the cells were passaged as small colonies with 0.5 mM EDTA (Gibco) in Ca^2+^/Mg^2+^-free DPBS (Gibco). The medium was supplemented with 5 μM of Y- 27632 (Selleckchem #S1049), a Rho-Associated Coil Kinase (ROCK) inhibitor, for the first 24 hours to prevent apoptosis after passaging. Cultures were tested for mycoplasma using a MycoAlert Kit (Lonza), and for bacteria, mold, fungi and yeast using solid and liquid lysogeny broth (LB, Sigma-Aldrich) and sabouraud dextrose agar (SDA, VWR) at 30°C and 37°C.

### Organoid differentiation

Cerebral brain organoids were differentiated according to a protocol published earlier^2^, with some modifications. Embryoid body differentiation was initiated on day 0 by detaching 70-80% confluent iPSCs with 0.5 mM EDTA and seeding 9 000 cells/well on ultra-low attachment U-bottom 96-well plates (CellCarrier-96 Spheroid ULA, Perkin Elmer 6055330) in 150μl of E8+0.5% P/S supplemented with 20 μM ROCK inhibitor. On days 1, 3 and 5, 120μl of medium was removed and 150μl fresh E8+0.5% P/S was added. Neuroinduction was mediated by a gradual transition from E8 to neuroinduction medium by replacing 150μl of the medium on days 6, 7, 8 and 10 to neurodinduction medium consisting of DMEM (Gibco #21331-020), 1x N2 (Gibco #17502-001), 1x Glutamax (Gibco #35050-038), 1x of nonessential amino acids (NEAA; Gibco #11140-035), 5U/ml of Heparin (LEO Pharma) and 0.5% P/S. On day 11, the spheroids were embedded to 20μl of matrigel and cultured in differentiation medium 1 (Diff1, 1:1 mixture of DMEM F12 and Neurobasal (Gibco #21103-049), 1x Glutamax, 0.5x NEAA, 0.5x of N2, 1x of B27 without vitamin A (Gibco #12587010), 2.5 μg/ml of insulin (Sigma #10516-5ml), 50 μM of 2-mercaptoethanol (Sigma) and 0.5% P/S). On d15, organoids were transferred onto 6-well plates (Sarstedt) in differentiation medium 2 (Diff2) with identical composition with Diff1, but supplemented with B27 containing vitamin A (Gibco; #17504001) instead of with B27 minus vitamin A (Gibco #12587010). From now on, organoids were maintained on orbital rotators (ThermoFisher Scientific #88881102) adjusted to 75 rpm. Medium was replenished every second day. To promote a good quality of organoids, the morphology of the spheroids was assessed under a microscope one-by-one before embedding to matrigel. The spheroids accepted for matrigel embedding were translucent, with some darkening of the core permitted, and maintained sharp edges. Common exclusion criteria were i) local expansion of tissue bulging out from the spheroid, i.e. not sharp edges ii) a foreign fiber translodged in the spheroid, iii) Incomplete formation of the embryoid body, where the well housed one main spheroid and one or more smaller spheroids attached to the bigger one. Also, on day 15, organoids without visible cortical loop structures were discarded. Finally, at any point on day 15+, organoids where the dense core had disintegrated, leaving a translucent piece of tissue, were also discarded.

### Differentiation of erythromyeloid progenitor cells

To initiate microglial differentiation on day 0, iPSCs were detached with 0.5 mM EDTA at 70-80% confluency, and 60 000-80 000 single cells/dish were seeded on 3.5 cm matrigel coated dish (Sarstedt) in mesodermal differentiation medium consisting of E8, 0.5% P/S, 5 ng/mL BMP4 (PeproTech), 25 ng/mL Activin A (PeproTech), 1 μM CHIR 99021 (Axon) and 10 μM ROCK inhibitor (Selleckchem #S1049). Cultures were maintained in a low oxygen incubator at 5% O2, 5% CO2 and 37°C. During the differentiation, medium was changed every 24 h. On day 1, the mesodermal medium was replaced but with lower 1 μM Y-27632. On day 2, within 44-46 h from seeding, hemogenic endothelial differentiation was induced by replacing medium with Dif-base medium consisting of DMEM/F-12, 0.5% P/S, 1% GlutaMAX, 0.0543% sodium bicarbonate (all from Thermo Fisher Scientific), 64 mg/L L-ascorbic acid and 14 μg/L sodium selenite (both from Sigma), and supplemented with 100 ng/mL FGF2, 50 ng/mL VEGF (both from PeproTech), 10 μM SB431542 (Selleckchem), and 5 μg/mL insulin (Sigma). On day 4, erythromyeloid differentiation was furthered by replacing the media by Dif-base supplemented with 5 μg/mL insulin, 50 ng/mL FGF2, vascular endothelial growth factor (VEGF), interleukin 6 (IL-6) and thrombopoietin (TPO), 10 ng/mL IL-3 and stem cell factor (SCF; all Peprotech). From now on, cells were maintained at atmospheric 19% O2, 5% CO2 and 37°C. On day 8, the erythromyeloid progenitors (EMPs) that were blooming into suspension were collected by gentle pipetting through a 100 μm strainer (VWR). The cells were counted and centrifuged at 160×g for 5 minutes to be ready for incorporation into organoids.

### Incorporation of microglial progenitor cells into organoids

To incorporate exogenous microglia into the organoids, day 8 microglial progenitors (EMPs) were allowed to spontaneously migrate into day 30 organoids using two comparable methods: either as free-floating cells or inside Matrigel droplet. For free-floating incorporation, single organoids were transferred to ultra-low attachment 96-well U-bottom wells (CellCarrier-96 Spheroid ULA, Perkin Elmer 6055330) in carryover medium. Then 500 000 EMPs/organoid, in 100μl of Diff2, were seeded on the wells. For control, Diff2 was added without progenitors. Organoids with (+)ORG and without (-)ORG microglia were incubated at 37°C, 5% CO2 for 6 hours to allow the progenitors to sediment and attach to the surface of the organoid. Then organoids were transferred back to 6-well format and the culture continued according to organoid maintenance protocol. For matrigel-droplet incorporation method, day 30 organoids were transferred to parafilm cups and day 8 microglial progenitors (EMPs) were resuspended in matrigel at 10 000 cells/μl density. Then 2 μl of the suspension was added on an organoid, giving 20 000 EMPs/organoid. The assemblies were then incubated at 37°C, 5% CO2 to allow the matrigel to solidify and settle on the organoids. The organoids were transferred to Diff2 the same way as on day 11 and the culture was then continued according to organoid maintenance protocol.

### Organoid fixation and immunohistochemistry

Organoids were washed thrice with 0.1 M phosphate buffer (REF) and fixed with freshly thawn 4% paraformaldehyde (Sigma) in phosphate buffer for 4 hours at 4°C. Fixed organoids were cryopreserved in 30% saccharose (VWR) in phosphate buffer at +4°C overnight. After which, they were embedded in O.C.T. Compound (Sakura, # 4583) in 1 x 1 x 0.5 cm plastic molds (Tissue-Tek) by incubating at RT for 20 minutes and freezing on a metal block at −70°C. Organoids were then cryosectioned to 20 μm thickness on cryotome (Leica CM1950), sections were mounted on microscope glasses (Thermo Scientific) and stored in −70°C. For heat-induced antigen retrieval, 10 mM sodium citrate (citrate (pH=6.0, VWR) was heated to 92°C, the slides were submerged and incubated for one hour, as the solution was allowed to cooled down in RT. Sections were blocked and permeabilized 10% normal goat serum (Millipore) and 0.05% Tween20 (Sigma) in PBS for 1 hour. Primary and secondary antibodies were diluted and incubated in the 5% normal goat serum in 0.05% Tween20 in PBS. Sections were incubated with primary antibodies at 4°C overnight, washed 3x in 0.05% Tween20 in PBS and incubated with species-specific AlexaFluor secondary antibodies (1:1000) for 1 hour at RT. Slides were mounted in Vectashield mounting medium with or without DAPI (Vector) for imaging. Staining controls that omitted the primary antibody were used to confirm staining specificity. All used antibodies are listed in Table 2.

**Table 2.** Antibodies used for immunocytochemistry.

| Antibody | Clone | Host | Pretreatment | Dilution | Cat. No | Producer |
| --- | --- | --- | --- | --- | --- | --- |
| IBA1 | - | RbP IgG | None | 1:500 | ab153696 | Abcam |
| IBA1 | - | RbP IgG | None | 1:250 | 01919741 | WAKO |
| CD41 | M148 | MM IgG2a | None | 1:1000 | ab11024 | Abcam |
| IBA1 | GT10312 | MM IgG1 | Na-citrate | 1:500 | MA5-27726 | Invitrogen |
| SOX2 | - | RbP | Triton | 1:200 | AB5603 | Millipore |
| TBR2 | WD1928 | MM IgG1κ | Triton | 1:100 | 14-4877-82 | eBioscience |
| DCX | - | RbP | Triton | 1:200 | 4604 | CST |
| SATB2 | - | RbP IgG | Na-citrate | 1:500 | ab34735 | Abcam |
| TUJ1 | TUJ1 | MM IgG2a, κ | Triton | 1:200 | 801202 | Biologend |
| PSD95 | D27E11 | RbM IgG | Na-citrate | 1:200 | 3450T | CST |
| SYN1 | SP11 | RbM IgG | Na-citrate | 1:250 | MA5-14532 | Invitrogen |

IgG, Immunoglobulin G; MM, mouse monoclonal; RbM, Rabbit monoclonal; RbP, rabbit polyclonal;

**Table 3.** IHC antibodies and Taqman primers used in characterising the MAD6 iPCS line

| Antibodies used for immunocytochemistry |  |  |  |
| --- | --- | --- | --- |
|  | Antibody | Dilution | Company Cat # and RRID |
| Pluripotency Markers | Mouse anti-Oct4 | 1:400 | EMD Millipore; cat. MAB4401 |
|  | Goat anti-Nanog | 1:100 | R&D Systems; cat. AF1997 |
|  | Mouse anti-SSEA4 | 1:400 | EMD Millipore; cat. MAB4304 |
|  | Mouse anti-TRA-1-81 | 1:200 | EMD Millipore; cat. MAB4381 |
| Differentiation Markers | SOX17 PE (Endoderm)<br>Otx2 AF488 (Ectoderm)<br>Brachyury APC (Mesoderm) | As per datasheet | R&D ICI9241P<br>R&D ICI979G<br>R&D IC2085A |
| Secondary antibodies | Goat anti-mouse Alexa Fluor 488 | 1:300<br>1:300<br>1:300 | Molecular Probes, cat. A11001<br>Molecular Probes; cat. A11004 |
|  | Goat anti-mouse Alexa Fluor 568<br>Donkey anti-goat Alexa Fluor 568 |  | Molecular Probes; cat. A11057 |
| **Primers** |  |  |  |
|  | **Target** | **Forward/Reverse primer (5'-3')** |  |
| **Primers** | **Target** | **Company** |  |
| Pluripotency Markers (qPCR) | *Nanog* | <u>Thermo Fisher Scientific; cat. Hs02387400 g1</u> |  |
|  | *Lin28* | <u>Thermo Fisher Scientific; cat. Hs00702808 s1</u> |  |
|  | *Sox2* | <u>Thermo Fisher Scientific; cat. Hs01053049 s1</u> |  |
| House-Keeping Genes (qPCR) | *ACTB* | <u>Thermo Fisher Scientific; cat. 4326315E</u> |  |
| Sendai virus | SeV | <u>Thermo Fisher Scientific; cat. Mr04269880_mr</u> |  |

### Confocal microscopy

All confocal images were obtained with a Zeiss Axio Observer inverted microscope (10x, 20x, 40x (oil) or 63x (oil) -objectives) equipped with LSM800 confocal module (Carl Zeiss Microimaging GmbH, Jena, Germany). DAPI and secondary antibodies were imaged with 405 nm (λex 353 nm/λem 465 nm), 488 nm (λex 495 nm/λem 519 nm) and 561 nm (λex 543 nm/λem 567 nm) lasers, respectively.

Imaging for quantification of PSD95 puncta and analysis of interaction with microglia:

For quantification of PSD95 puncta, we manually sampled four optical fields per organoid, four organoids per (+)/(-)ORG group, and captured a Z-stack of ten 109.38 μm x 109.38μm (1580 px x 1580 px) focus planes with 0.28 μm interval, using the Plan-Apochromat 63x/1.40 oil DIC M27 objective. We applied the same imaging parameters for assessing interactions between microglia and synapses, with maximum orthogonal projections acquired at ZEN 2.3 Blue edition or Zen 3.2 lite edition for the representative images in Figure 2.

For quantification of morphology, we employed two imaging devices to acquire whole slice Z-stack tiles, capturing the entire Z-dimensional morphology of the microglia. We imaged the day 120 organoids at the Zeiss confocal, with seven focus planes (merged into maximal orthogonal projection), using the EC Plan-Neofluar 10x/0.30 M27 objective. Day 35 and day 66 organoids we imaged using 3D HISTECH Pannoramic 250 Flash III with 20x objective (3DHISTECH Ltd, Budapest, Hungary) and the extended view images exported to TIF for downstream analysis using the Caseviever version 2.4.

For imaging of representative figures we used Leica Thunder 3D tissue imager, Fig. 1a, (Leica microsystems CMS GmbH, Wetzlar, Germany), 10X objective and six focus planes and created the orthogonal projection using the Leica X application suite version 3.7.2.22383. The Zeiss confocal with 20x objective for Fig. 1c-f, 40x for Fig. 1 g and 63x for Fig. 1i.

### Quantification of immunohistochemistry

The analysis on microglia morphology was done with the original CZI files (Zeiss LSM800 confocal), or the TIF files (3D HISTECH Pannoramic 250 Flash III). Skeletal analysis was performed using Fiji^42,43^. For the AI-based detection, the images were further compressed to a wavelet file format (Enhanced Compressed Wavelet, ECW, ER Mapper, Intergraph, Atlanta, GA) with a target compression ratio of 1:5. The compressed virtual slides were uploaded to a whole-slide image management server (Aiforia Technologies Oy, Helsinki, Finland). A convolutional neural network-based algorithm was trained to detect ramified, intermediate, rod and spherica cells using cloud-based software (Aiforia version 4.8, Aiforia Create, Aiforia Technologies Oy, Helsinki, Finland). We performed counting of the PSD95 puncta with a set of 12 prominence thresholds, using the Find Maxima algorithm in Fiji^42^. Final results were obtained via Origin 2019b (OriginLab, Northampton, Massachusetts, USA).

### Pro-inflammatory stimulation cerebral organoids

To study proinflammatory stimulation on organoids, a single organoid per well was transferred on 6 well plates on day 77 and 120 in Diff2 medium with 20 ng/ml LPS serotype O111:B4 (Sigma # L2630) and incubated for 24 h. After incubation, the medium was centrifuged at 300×g for 5 minutes to remove cell debrisand stored at −20 °C until analysis for cytokine measurement. Organoids were fixed for immunohistochemistry.

### Cytokine secretion

Quantitative cytokine concentration was determined using a Cytometric Bead Array (CBA) with multiplexing cytokine flex sets (BD Biosciences). For CBA, 20 μl of samples, blanks and cytokine standards were incubated with equal volumes of bead mixture (with final bead dilution of 1:75) and then detection reagent (1:75). Human Soluble Protein buffer kit (BD Biosciences) was used in conjunction with flex sets for detecting granulocyte-macrophage colony-stimulating factor (GM-CSF), interleukins IL1a, IL1b, IL6, IL8 and IL10, monocyte chemoattractant protein-1 (MCP1, CCL2), regulated on activation normal T cell expressed and secreted (RANTES, CCL5), and tumor necrosis factor alpha (TNFa; all BD Biosciences). The samples were run on a Cytoflex S (Beckman Coulter, California, USA) and a minimum of 300 events for each cytokine was measured. Channels 638/660 nm (APC) and 638/780 nm (APC-A750) excited by a 638 nm red laser were used for determining the positions of bead clusters. Cytokine level was measured with PE-reporter measured via 561/585 nm channel exited with a 561 nm laser. The data were analyzed on FCAP Array™ v2.0.2 software (BD Biosciences) and cytokine concentrations were calculated with quantitative logarithmic 4-parameter analysis from a standard curve.

### Brain slice preparation for electrophysiology

Brain organoids (107-165 days in culture) were embedded in 2% low melting point agarose and sliced (for whole-cell recordings, 350 μm; for multielectrode recordings, 500 μm) using a vibratome (Campden Instruments, model 7000 smz) in chilled (4 °C), fully carbonated (95%/5%, O2/CO2) aCSF of the following composition (in mM). 92 N-methyl-D-Glucamine, 2.5 KCl, 20 HEPES, 25 NaHCO3, 1.25 NaH2PO4, 3 Na-Pyruvate, 2 Thiourea, 5 Na-Ascorbate, 7 MgCl2, 0.5 CaCl2, 25 Glucose (pH adjusted to 7.3 with HCl 10 M). After cutting, slices were placed in a custom-made chamber and allowed first to recover at 34 °C for 30 minutes in the recording solution (see below for composition, supplemented with 3 Na-Pyruvate, 2 Thiourea and 5 Na-Ascorbate) and then for 60 minutes in the same solution at room temperature (20-22°C) before use for electrophysiology.

### Whole-cell electrophysiology

A single slice (350 μm) was transferred and secured with a slice anchor in a large volume bath under an Olympus BX50WI microscope equipped with differential interference contrast (DIC) optics, an x40 water immersion objective and a charge-coupled device (CCD) camera (Retiga R1, Q-imaging). Slices were continuously perfused at a rate of 2.5-3 ml/min with a recording solution of the following composition (in mM): 120 NaCl, 2.5 KCl, 25 NaHCO3, 1.25 NaH2PO4, 2 CaCl2, 1 MgCl2, 25 Glucose. Organoids from both phenotypes were examined on a single experimental day with a blind protocol to the experimenter. Whole-cell current and voltage-clamp recordings were conducted at 32-23°C with an Axopatch-200B amplifier (Molecular Devices) using 5-8 MOhm glass electrodes filled with an internal solution containing (in mM) 135 Potassium-gluconate, 5 NaCl, 10 HEPES, 2 MgCl2, 1 EGTA, 2 Mg-ATP, 0.25 Na-GTP & 0.5% Biocytin (pH adjusted to 7.3 with osmolarity at 275-285 mOsm/l). Electrophysiological data were low pass filtered at 1 KHz (4-pole Bessel filter) then captured at 10 KHz via a Digidata 1440A A/D board to a personal computer, displayed in Clampex software (version 10.7, Molecular Devices) and stored to disk for further analysis. Single-cell electrophysiological current and voltage-clamp data were analyzed in Clampfit (Molecular Devices). Synaptic data were detected and analysed with Mini Analysis software (Synaptosoft Inc).

### 3D-Multielectrode electrophysiology

A single slice (500 μm) was transferred and secured with a slice anchor in the chamber of a 60-3D- multielectrode recording array (3D-MEA, Multichannel Systems-MCS) with a 8×8 configuration (electrode impedance, 100-200 KOhm; spacing, 250 μm; height, 100 μm, conducting area only on top 20 μm) made of titanium nitride (TiN). Recordings were made with a MEA2100-Mini headstage (MCS, Germany) under a Leica S APO microscope equipped with a transillumination base and a digital camera (model MC170HD, Leica Microsystems Germany). Slices were continuously perfused at a rate of 3-3.5 ml/min at 32-23°C with the same recording solution as with whole-cell recordings and were allowed at least 20 minutes to settle in the MEA before any recordings or pharmacology was attempted. Organoids from both phenotypes were examined on a single experimental day with a blind protocol to the experimenter. Electrophysiological data were band passed between 300 and 3000Hz (second order Butterworth filter) and were captured at 20 KHz via MCS-IFB 3.0 multiboot (MCS, Germany) to a personal computer, displayed in MCS Experimenter software (version 2.15) and stored to disk for further analysis. Multielectrode recording data files were first converted and then imported into Spike2 (Cambridge Electronic Design, UK) or Neuroexplorer (Nex Technologies, US) software for measurements and spike sorting.

### Statistical analysis

All data represent mean±s.e.m. The morphologies of microglia were analysed on Origin (OriginLab Corporation, Northampton, USA) with significance tested using the Tukey test. All other statistical tests were performed on Prism (GraphPad Software, La Jolla California USA), with two-way-ANOVA for the cytokine measurements, and non-parametric Mann-Whitney unpaired tests for PSD95 puncta and electrophysiology results (except for paired T-test between (-)ORG & (-)ORG + NMDA in Fig. 4h).

## Acknowledgements

This work was supported by grants from the Academy of Finland (328287, 305516 and 301234) and the Finnish brain foundation. We also thank Mrs Mirka Tikkanen, Antonie Mugnier and Shaila Eamen for their technical help, and Jenni Voutilainen, Ida Hyötyläinen and Sara Wojciechowski for assistance characterising the generated Mad6 iPSC line and Markku Laakso for providing the original fibroblasts. This work was carried out with the support of UEF Cell and Tissue Imaging Unit, part of Biocenter Finland, University of Eastern Finland, Finland, and FIMM Digital Microscopy and Molecular Pathology Unit supported by HiLIFE and Biocenter Finland.

## Author contributions

IF and AD designed the work, conducted the experiments and interpreted the results with input from AS, M G-B, PK, HK, SO and FF in the experimental work. MK trained us to culture cerebral organoids. AP, DH, JK and SL provided the iPCS lines.TM directed the project, interpreted data and supervised the work. IF, AD, HK and TM wrote the manuscript and all co-authors read and commented the manuscript and approved the final version

## Data availability

All relevant data is available from the authors upon reasonable request.

## Supplements

**Supplement Figure 1.**
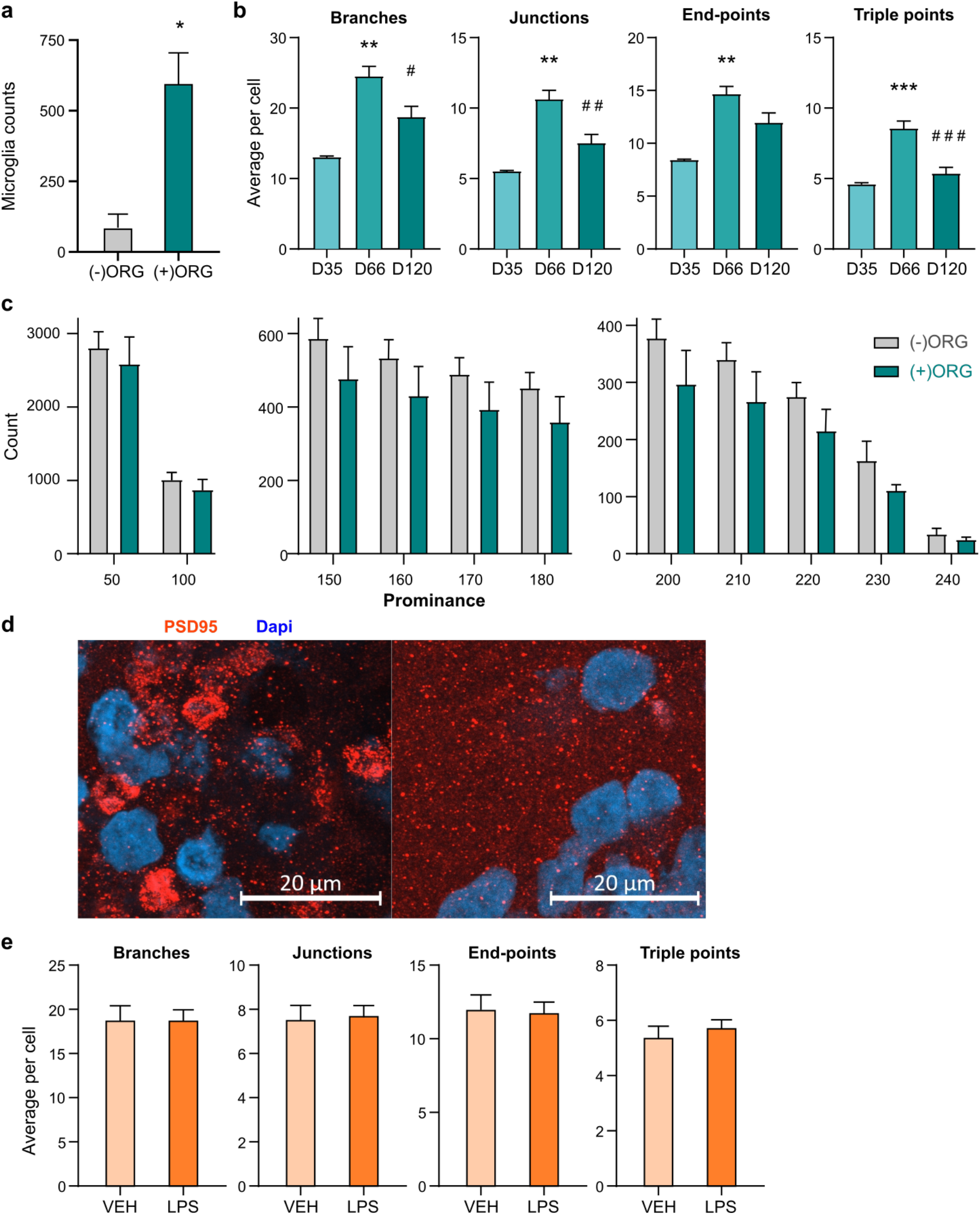
**a** The sections from (+)ORGs hold significantly more Iba+ cells than from (-)ORGs. The data bars present means (+/- standard error of mean) from five (-)ORGs against 18 (+)ORGs. **b** Skeletal analysis of Iba1+ cells in organoids at days 35, 66 and 120 showed a significant increase in complexity from day 35 to 66 but decreased from day 66 to 120. The bars represent means (+/- SEM) from nine VEH organoids against ten LPS exposed organoids. Significance ***p < 0.001, **p < 0.01, *p < 0.05 compared to D35, # indicate similarly significance compared to D66; **c** Skeletal analysis of Iba1+ cells in organoids exposed to 24h LPS or vehicle treatment. The bars represent the mean (+/- SEM) percentage of three organoids for the day 35 group, six for day 66 and nine for day 120. The bars represent means (+/- SEM) from nine VEH organoids against ten LPS organoids. **d** Counts of PSD95 puncta in orthogonal projection images of PSD95 immunohistochemical stainings, presented in a gradient of the increasing puncta prominence. The bars present means (+/- SEM) of a total of 16 identically acquired confocal orthogonal projection images of immunohistochemical stainings against PSD95. N = 4 organoids. **e** Immunohistochemical stainings of day 120 cerebral organoid cryosections show PSD95 localization as scattered puncta in post-synaptic termini, and in cell soma.

**Supplement Figure 2.**
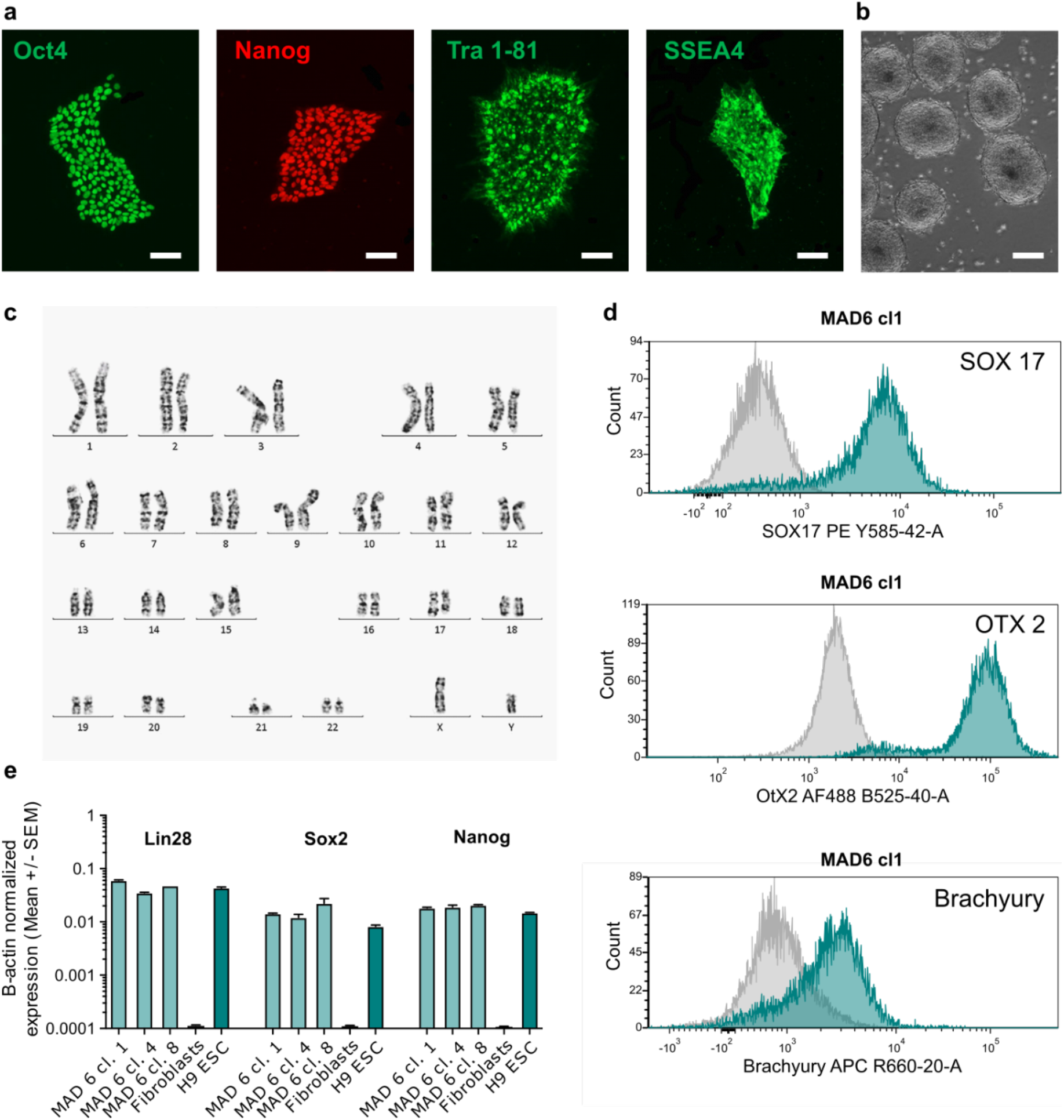
**a** IHC detection of the pluripotency markers of Oct4, Nanog, Tra 1-81 and SSEA4 expressed in iPCS colonies. Scale bars 100μm. **b** Generation of embryoid bodies to test for differentiation to all three embryonic layers. Scale bar 50μm. **c** Karyotyping results showed normal 46 XY karyotype. **d** Detection of markers expressed in endodermal (SOX17), ectodermal (OtX2) and mesodermal (Brachuyry) lineages, measured in flowcytometry analysis of single cell suspension of the embryoid bodies. **e** RT-qPCR measurements of the expression of the pluripotency markers Lin28, Sox2 and Nanog across three clones of MAD6, with fibroblasts and an embryonic stem cell line (H9 ESC) as negative and positive controls, respectively.

